# Understanding the Divergent Evolution and Epidemiology of H3N8 Influenza Viruses in Dogs and Horses

**DOI:** 10.1101/2023.03.22.533763

**Authors:** Brian R. Wasik, Evin Rothschild, Ian E.H. Voorhees, Stephanie E. Reedy, Pablo R. Murcia, Nicola Pusterla, Thomas M. Chambers, Laura B. Goodman, Edward C. Holmes, James C. Kile, Colin R. Parrish

## Abstract

Cross-species virus transmission events can lead to dire public health emergencies in the form of epidemics and pandemics. One example in animals is the emergence of the H3N8 equine influenza virus (EIV), first isolated in 1963 in Miami, Florida, USA, after emerging among horses in South America. In the early 21^st^ century the American lineage of EIV diverged into two ‘Florida’ clades that persist today, while an EIV transferred to dogs around 1999 and gave rise to the H3N8 canine influenza virus (CIV), first reported in 2004. Here, we compare CIV in dogs and EIV in horses to reveal their host-specific evolution, to determine the sources and connections between significant outbreaks, and to gain insight into the factors controlling their different evolutionary fates. H3N8 CIV only circulated in North America, was geographically restricted after the first few years, and went extinct in 2016. Of the two EIV Florida clades, clade 1 circulates widely and shows frequent transfers between the USA and South America, Europe and elsewhere, while clade 2 was globally distributed early after it emerged, but since about 2018 has only been detected in Central Asia. Any potential zoonotic threat of these viruses to humans can only be determined with an understanding of its natural history and evolution. Our comparative analysis of these three viral lineages reveals distinct patterns and rates of sequence variation yet with similar overall evolution between clades, suggesting epidemiological intervention strategies for possible eradication of H3N8 EIV. (242 words)

**IMPORTANCE:** The emergence of viruses in new hosts is a threat to human and animal health. The H3N8 equine influenza virus (EIV) emerged in 1963 by transfer of an avian influenza virus, and the H3N8 canine influenza virus (CIV) subsequently emerged in 1999 when EIV transferred to dogs. H3N8 CIV persistently circulated in only a few locations in the USA, and has not been detected since 2016. In the same period H3N8 EIV has circulated as two separate clades, one in North America and other regions of the world, while the other currently appears to be found only in Central Asia. By comparing the hosts, epidemiology, and evolution of these influenza viruses we explain how these lineages had different evolutionary fates, and show why elucidating these evolutionary processes is key to understanding zoonotic disease and viral emergence. (137 words)

## Introduction

The processes driving virus outbreak epidemiology and cross-species virus transmission are topics of great importance. Studying viruses involved in emerging outbreaks in animals can reveal the fundamental processes that mediate successful transfer between hosts and the subsequent establishment of viral variants as epizootic or even zoonotic and pandemic pathogens. Such studies also suggest approaches to protecting humans and other animals from potential zoonotic viruses by preventing cross-species epizootic outbreaks, as well as new methods for blocking the emergence of transferred viruses or reducing the adaptation of viruses to new hosts, thereby reducing their pandemic potential (1–4).

Most subtypes of influenza A virus (IAV) are well established in their reservoir hosts, mainly birds living in aquatic environments (5). However, several subtypes appear to have a relatively high propensity for infecting other animal hosts, including domestic poultry, humans, swine, ferrets, seals, horses, and most recently dogs (5, 6). Some mammals occasionally suffer outbreaks from avian origin influenza, including seals and mink notably during recent spread of highly pathogenic avian influenza (HPAI) H5N1, that may suggest new subtype risks (7, 8). Most spillovers cause short-term outbreaks that rapidly die out, while some have transitioned from emergent to epidemic or pandemic pathogens in their mammalian hosts. In human populations, some IAV subtypes (primarily H1N1 and H3N2 in the modern era) have become endemic diseases (9). IAV are members of the family *Orthyomyxoviridae*, and have a negative-sense RNA genome arranged as eight segments: PB2, PB1, PA, HA, NP, NA, M, and NS (9).

Horses have historically been an important mammalian host for IAV, likely including epidemics in the 1870s in North America that posed a significant burden on animal health, as well as on a human society that was highly dependent on horses (10). In the 20^th^ century, horses were infected by an H7N7 virus that was first recognized in 1956 and became extinct in the late 1970s (11). H3N8 equine influenza virus (EIV) was first isolated in 1963 in Florida, likely after being introduced along with infected horses from South America, and descendants of that virus continue to circulate in horses today (12–15). The appearance and circulation of reassortant H7N7 and H3N8 EIVs in the 1970s suggests that the H3N8 contributed to displacement and extinction of the H7N7 subtype. Vaccines against H3N8 EIV were first introduced in the late 1960s, and the period of the 1970s and early 1980s resulted in several subclade divergences and increasing H3N8 diversity (13, 16). Around the year 2000 the H3N8 EIV population divided into different lineages: the Florida Clade 1 (FC1) was primarily in North America but also observed in outbreak in other regions of the world, while the Florida Clade 2 (FC2) was found primarily in Europe, yet both virus clades have spread to many different countries and continents (17–22). However, these clades appear to have kept some degree of geographic separation, and they do not appear to have been in direct competition (23).

EIV is a common respiratory tract infection in horses and closely related equids, causing influenza-like signs similar to those seen in humans (24). Current estimations of EIV incidence in US horses comes from voluntary surveillance, and are frequently under 10% with seasonal variability (25, 26). Vaccines for EIV have been available since the 1970s and include antigens from the most commonly circulating strains, and those are widely used in domesticated horses (27). While human influenza vaccines are updated frequently with the most widespread antigenic variants, EIV vaccines are updated less often. In the United States, roughly 60% of equine operations vaccinate for EIV (66.4% in 1998, 54.1% in 2005, 60.4% in 2015) (28). It is unclear to what degree low vaccine effectiveness or antigenic mismatch have contributed to recently observed ‘breakthrough’ outbreaks in multiple countries (22, 29–32).

Recommendations for vaccine formulations are regularly proposed by the World Organization for Animal Health (formerly Office International des Epizooties, OIE) expert surveillance panel (33, 34), but strain inclusion varies in different vaccine formulations (35). The potential for horses to be sources of zoonotic disease given their close association with humans makes EIV an identified neglected disease threat of concern (36, 37). Experimental inoculation of human volunteers in the 1960s showed that the virus was able to replicate in humans with sufficient exposure (38, 39). Only through proper surveillance and sequencing of circulating EIV in different populations will a clearer picture of the virus variation be obtained. In particular, this would enable targeted antigenic testing to justify the production of vaccines with a better antigenic match and greater protective efficacy (34, 40). Determining the relationships between viruses causing different outbreaks through surveillance will also reveal epidemiological pathways that arise during epidemics, allowing better strategies for control and potentially eradication.

A single H3N8 EIV transferred to dogs around 1999 to create the first known epidemic of canine influenza virus (CIV). Molecular and epidemiological evidence suggests the epidemic originated from a single, cross-species transmission event of a non-reassortant H3N8 EIV, with cases being initially identified in greyhounds in a training facility in Florida in 2003 and 2004 (41–44). The H3N8 CIV spread to several regions of the USA around this period, likely due to transport of infected greyhounds, although the virus only achieved prolonged transmission in dogs within larger dense populations, including animal shelters, kennels, and canine day cares (45). The virus appears to have persisted in large metropolitan areas (e.g. Denver and Colorado Springs, New York City, and Philadelphia) with large shelter dog populations that had high standing populations and high rates of turnover (46). While the emergence of CIV H3N8 led to an epidemic, other observed spillovers of EIV H3N8 into dogs (in 2002 in the UK and 2007 in Australia) did not result in onward canine transmission (47, 48). No human infections with H3N8 CIV were observed, despite frequent interactions (49). A second CIV arose around 2005 in Asia, and that H3N2 strain appears to have been derived directly from an avian influenza virus that transferred to dogs (50). H3N2 CIV appears to have circulated widely in Korea and China for a number of years, and has been repeatedly introduced into the USA and Canada where it has mostly caused local and self-limiting outbreaks (51). Risk assessment of H3N2 CIV transmission to humans was found to be minimal (52).

Host transfers underpin many infectious disease epidemics or pandemics, and in recent years such events have given rise to new emerging viruses in humans and other animals, including the 2009 pandemic swine influenza (pdmH1N1), Ebolavirus, SARS-CoV-1 and -2, and the H3N8 and H3N2 CIV (1, 53). Influenza surveillance networks (such as the World Health Organization Global Influenza Surveillance & Response System, WHO GISRS) have made considerable leaps in the past decade, with rapid identification of human cases of highly pathogenic avian influenza (54, 55). Despite this, gaps remain in our ability to predict changes in host-range that result in virus emergence, or to create early warning systems to detect and mitigate outbreaks before they get underway (4, 56, 57). Strategies for controlling emerging viruses primarily involve aggressively managing outbreaks after they first occur. Yet, this requires a clear understanding of the routes of transmission, ongoing circulation, and likely host adaptation leading to increased spread or immune evasion, and such knowledge is often lacking in the early stages of outbreaks (3). It is also often unclear whether and how virus evolution in the original host facilitates the initial successful spillover infection, or how the viruses later evolve and adapt to their new hosts compared to evolution that occurs in their original hosts. Understanding these processes in more detail may allow us to anticipate and forestall the emergence of other viruses in the future.

As well as local outbreaks, a defining feature of the most threatening virus emergence events is longer distance and international transfers that allow viral outbreaks to sustain over time and develop into pandemics, particularly within human host populations (58, 59). Long-distance transfers are also a feature of H3N8 EIV epidemiology, likely associated with the movement of infected racing and sport horses (31, 60, 61). It is estimated that just 0.7% of operations transport horses outside North America, yet these international transfers of horses have made EIV a global pathogen of concern (28, 34, 62). Among dogs, H3N2 CIV has similarly transferred between continents (51). In contrast, the H3N8 CIV outbreaks were more confined, circulating within and between animal shelters and kennels within the central and eastern regions in the United States (45, 46). Notably, that virus lineage did not appear to spread outside the USA.

Modern genomic epidemiology has allowed the detailed examination of viral outbreaks, including the spread and replacement of viral variants (2, 63–65). For example, the emergence of more transmissible or antigenic variant viruses that displace earlier lineages is commonly seen in evolving virus populations, including 2009 pdmH1N1, SARS-CoV-2 (B.1.1.7 ‘Alpha’, or B.1.1.529 ‘Omicron’), and the canine parvovirus type-2a variant that emerged and efficiently replaced the earlier strains of that virus (66–72). These events highlight the importance of continued surveillance and analysis of viruses; both in new hosts and in their original hosts, as such comparative analysis allows us to identify mechanisms of true adaptive evolution following emergence.

To resolve the evolution of H3N8 in dogs and horses we collected genomic data from our own sequencing and public repositories of H3N8 beginning around 2001. Complete genome sequences of all segments enable phylogenetic relationships and host-specific evolution to be determined. We further employed use of the much larger collection of HA1 sequences (HA1 being the hemagglutinin ‘head’ containing the receptor binding domain and dominant antigenic epitopes, H3 numbering 1-329). Here, we investigate the emergence the H3N8 canine influenza virus (CIV) and its origin from the H3N8 equine influenza virus (EIV), describe the later extinction of H3N8 in dogs while also following the continuing evolution and epidemiology of EIV among horses in several geographic regions of the world.

## Results

### H3N8 experienced major clade divergence in the early 2000s, including the emergence of CIV in dogs

Despite years of spread and evolution in horses since its first emergence around 1963, H3N8 EIV diversity had largely collapsed to a single North American clade by the end of the 20^th^ century (Figure 1, inset) (17, 73). This ‘Florida’ clade then gave rise to the FC1 and FC2 subclades. A full genome phylogeny since 2001 shows the presence of pre-clade divergence viruses (preC), and viruses within the monophyletic FC2, FC1, and CIV groups. Divergence occurred roughly after the appearance of Kentucky/5/2002, California/8560/2002, and Newmarket/5/2003 preC (Figure 1), so that the first full-genome samples from the FC2, FC1 and CIV clades are Richmond/1/2007, Wisconsin/1/2003, ca/Florida/242/2003, respectively. These FC2 and FC1 sequences are currently (or closely match) the EIV formulations frequently manufactured and used for vaccination (35).

**Figure 1.**
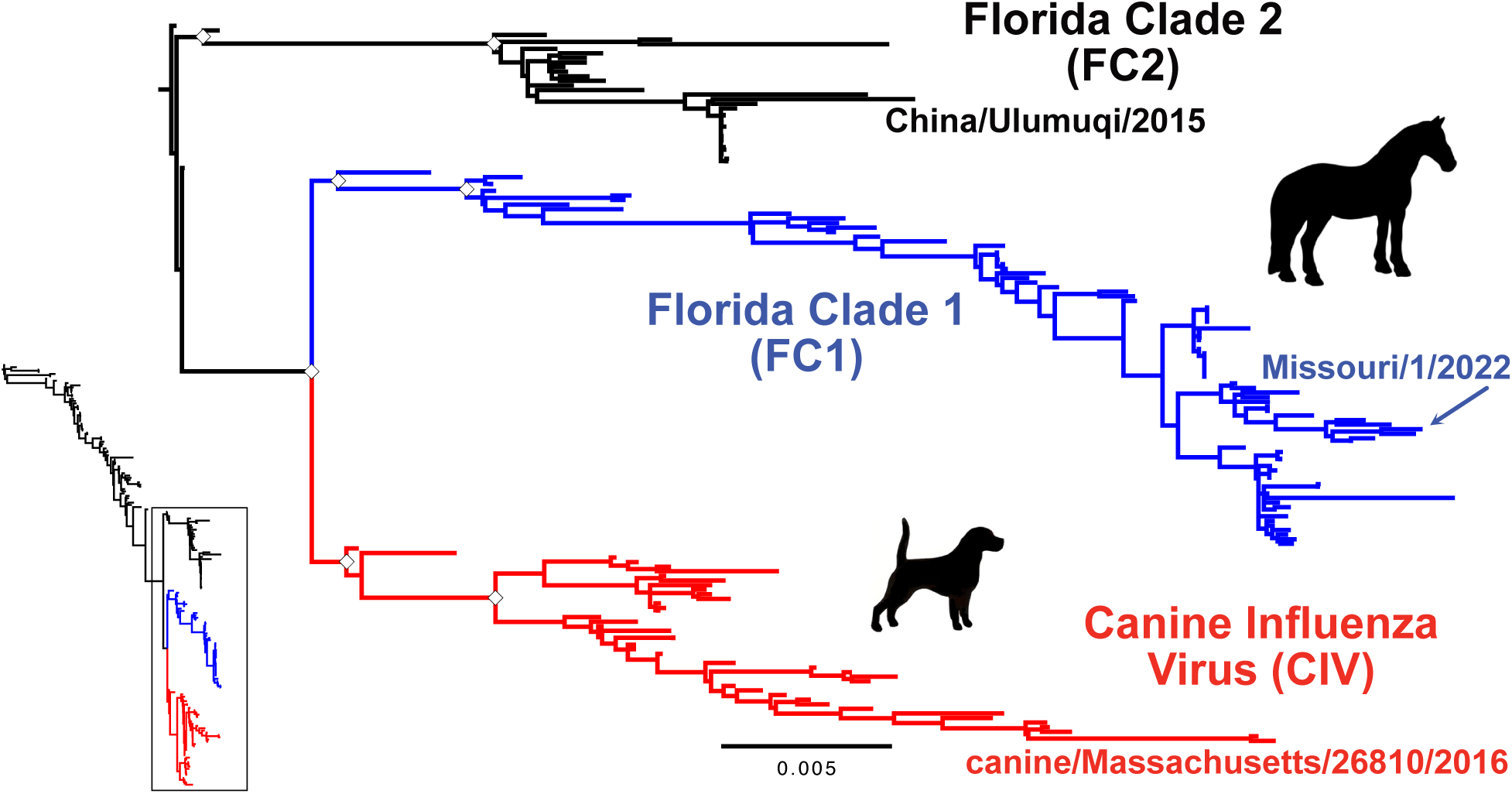
Recent evolution of H3N8 influenza virus. Equine influenza virus (EIV) H3N8 evolution from 1963 to present has resulted in three distinct clades post 2002 (inset). ML tree of the whole genome data set revealing three distinct clades: the Florida Clade 1 (FC1) and Florida Clade 2 (FC2) in horses, and Canine Influenza Virus (CIV) in dogs. Tree was rooted to pre-clade (preC) genome Kentucky/5/2002. Scale denotes nucleotide divergence. White diamonds at major early nodes represent bootstrap support >99%. Most recent genome of each clade is noted.

We further gathered all detailed collection dates at best accuracy (YYYY-MM-DD) to review the temporal nature of the data set. A root-to-tip regression analysis of the full genome data set revealed strong clock-like structure (R^2^ = 0.92) and an evolutionary rate of 1.5 x 10^-3^ nucleotide substitutions/site/year (Figure S1A). It was obtained a reliable root-to-tip regression for the expanded HA1 data set from 2001 to present, yet with rooting divergence from the common ancestor between FC1/CIV and FC2 lineages (Figure S1B). The establishment and persistence of these lineages allows a direct comparison of H3N8 influenza virus evolution in novel (dog) and original (horse) hosts following the host transfer and emergence event giving rise to CIV. To determine what genomic characteristics may have allowed FC1 (and FC2) to persist in nature, while CIV went extinct, we analyzed the CIV, FC1 and FC2 clades since their divergence in parallel (Figure 1).

### CIV was restricted to the United States and went extinct around 2016

The last known positive cases of H3N8 CIV were reported in the Northeast USA in 2016, two samples of which (Maine and Massachusetts) were sequenced shortly thereafter by our group. No further H3N8 positive cases in the USA have been reported, even though this was a time of frequent canine influenza testing due to the emergence and spread of H3N2 CIV in the US in 2015 and 2016 (51).

We gathered a data set of 43 complete CIV genomes (and an expanded 88 HA1 reading frame sequences) and employed the same methods as for the full H3N8 data set. Our full genome CIV phylogeny (Figure 2A) closely matches the monophyletic group seen in the larger H3N8 full genome data set (Figure 1) and previous analyses (74). The full genomes of CIV showed a strong clock-like evolution (R^2^ = 0.94) at approximately the same rate (1.8 x 10^-3^ substitutions/site/year, Figure S1C) as observed with the larger H3N8 full genome data set. We used the expanded HA1 data set to connect the phylogenetic tree to the geographic data (Figure 2B). Subclades of CIV were highly structured by regional introduction and spread, mostly localizing to outbreak centers in Florida, New York City, Philadelphia, and the cities of Colorado. Secondary geographic sites appear to have involved secondary outbreaks that were related by phylogenetics back to their source locations.

**Figure 2.**
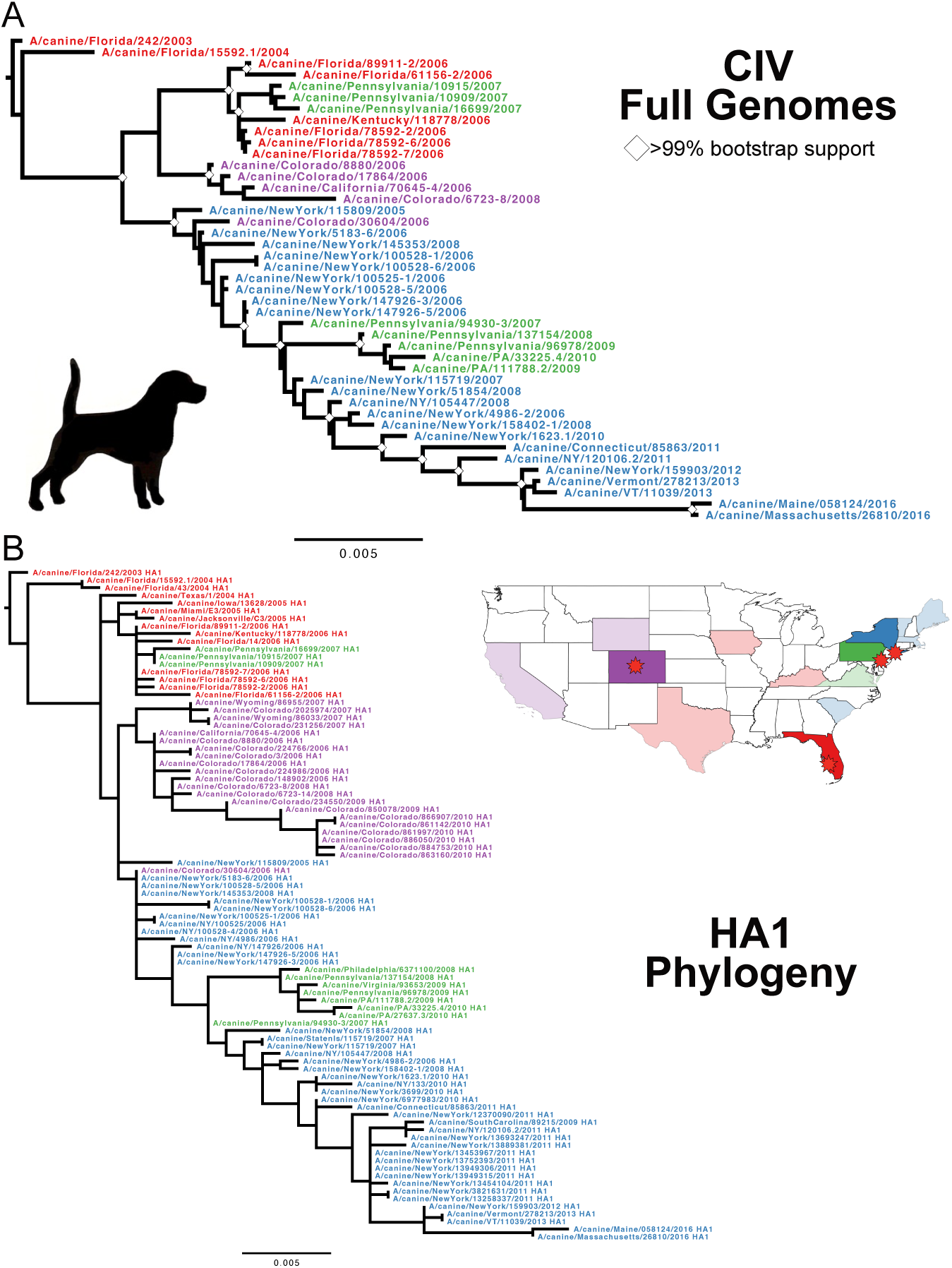
Evolution of the CIV H3N8 clade. (A) ML tree of the full genome data set of CIV, with taxa color defined by outbreak source region: Red, Florida/SE; Blue, New York City; Green, Philadelphia; Purple, Colorado. Tree was rooted to earliest sample, Florida/242/2003. Scale denotes nucleotide divergence. White diamonds at major nodes represent bootstrap support >99%. (B) ML tree of expanded HA1 data set reveals the strong geographic organization of subclades.

We further inferred the demographic history of these populations using Bayesian analysis with a Skyline Plot (BSP) tree prior, enabling us to estimate changes in genetic diversity – a measure of effective population size (Ne) – through time. For both data sets we found that Ne plots correspond to virus natural history and epidemiology: large population and diversity growth occurring during the initial US-wide outbreaks of 2005-2007, followed by regional die out and the eventual complete extinction (Figures S2A-B).

### EIV FC1 is a US-centric virus that contributes to international transfers and outbreaks

We compiled 82 FC1 full genome and 252 HA1 sequence data sets including those newly generated here. The FC1 full genome phylogeny shows a fairly linear clade divergence (Figure 3). The trees show strong signatures of geographic clustering, suggesting frequent single-introduction outbreaks in particular locations including South America, Europe, and other regions. There is a strong clock-like evolution (R^2^ = 0.99) at approximately the same rate (1.7 x 10^-3^ substitutions/site/year, Figure S1E) seen in the larger H3N8 full genome data set and the CIV clade. We again estimated Ne changes through time, and while sampling bias and poor admixture limits making broad conclusions, we were able to identify key demographic transitions and data gaps (Figures S2A, S6, S7).

**Figure 3.**
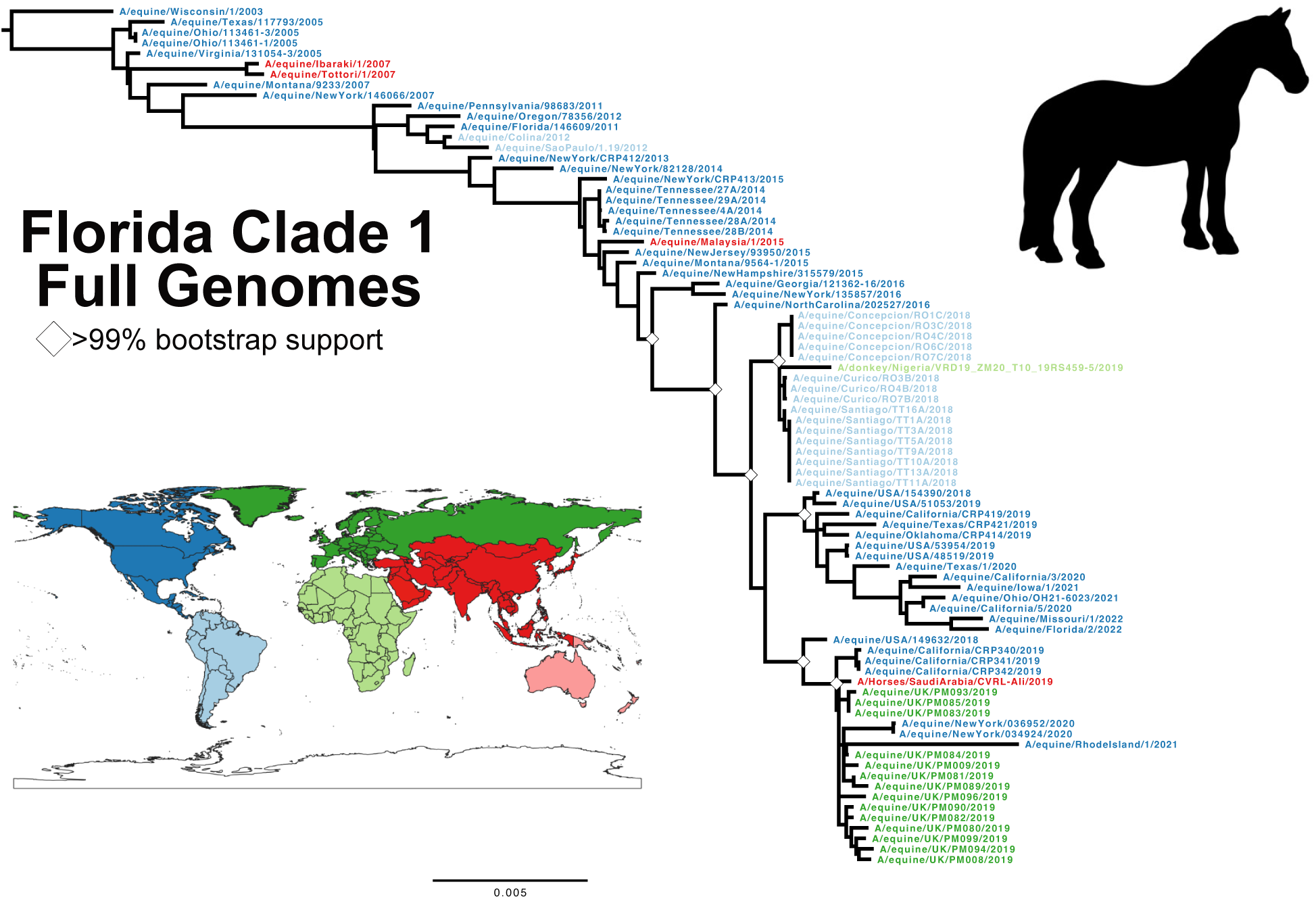
Evolution of the H3N8 EIV clade FC1. ML tree of full genome data set of FC1, with taxa color defined by worldwide region. Tree was rooted to earliest sample, Wisconsin/1/2003. Scale denotes nucleotide divergence. White diamonds at major nodes represent bootstrap support >99%.

The phylogenetic tree of our expanded HA1 sequences largely paralleled the full-length genome tree, further clarifying the larger geographic movements of FC1 since its divergence. The overall phylogenetic structure reveals an apparent US-based continuous circulation that seeds multiple international transfers and outbreaks in different regions of the world (Figure 4). Some of these historical transfers have been reported and analyzed previously and confirmed here (20, 75–79). Our Bayesian demographic analysis of the FC1 HA1 data set (Figure S8) also shows that the effective population size largely mirrors the full-length genome data set, yet with more points of transition and changes in diversity, some of which are likely revealed by the increased sampling resolution (Figure S2B).

**Figure 4.**
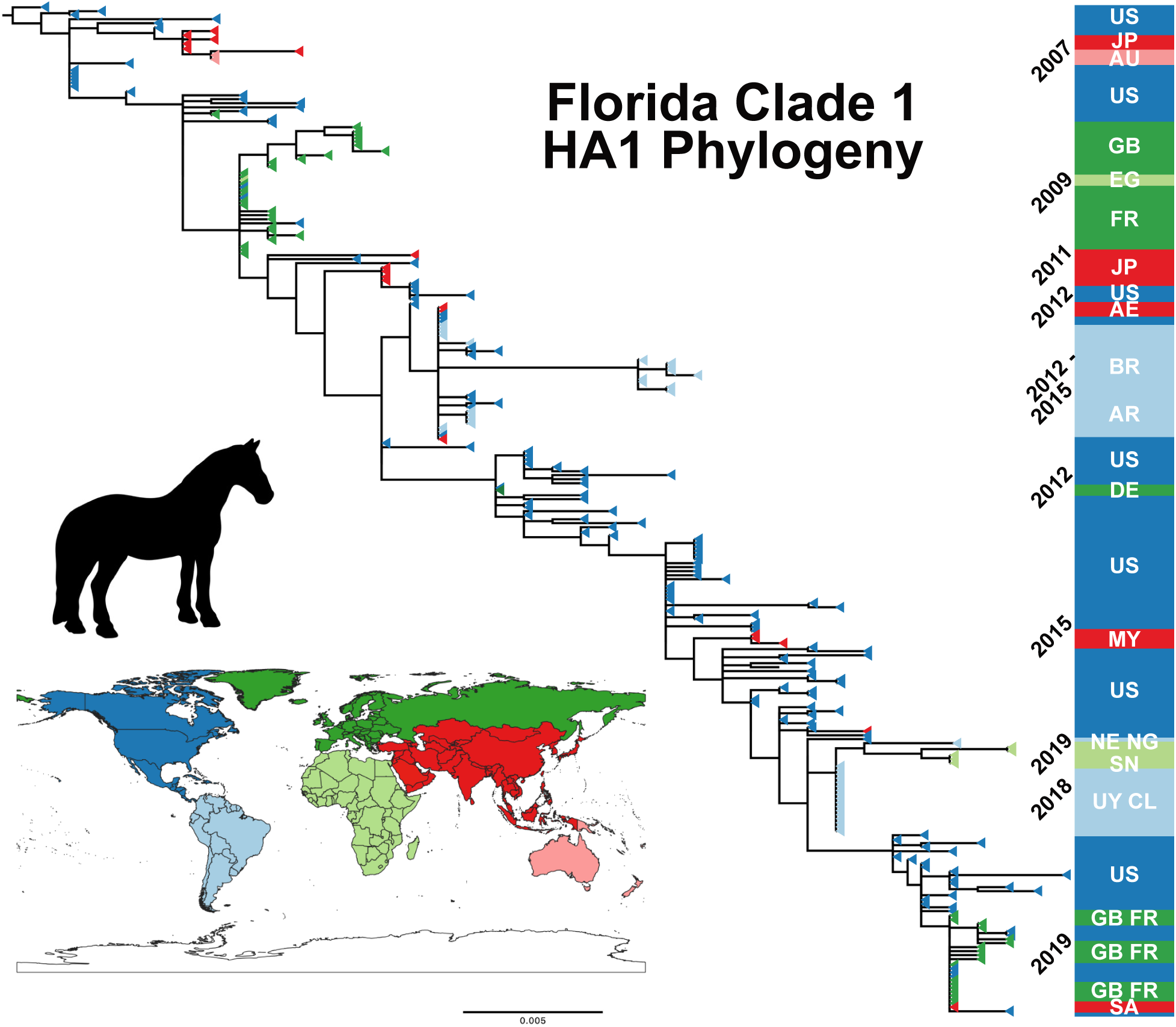
Evolution of the H3N8 EIV clade FC1. ML tree of the expanded HA1 data set (n=251), also colored by regional isolate. Tree was rooted to earliest sample, Wisconsin/1/2003. Scale denotes nucleotide divergence. Dates of and geography of regional subclades shows the recurring impact of US circulation on international transfers. Country ISO codes used: AE, United Arab Emirates; AR, Argentina; AU, Australia; BR, Brazil; CL, Chile; DE, Germany; EG, Egypt; FR, France; GB, United Kingdom; JP, Japan; MY, Malaysia; NE, Niger; NG, Nigeria; SA, Saudi Arabia; SN, Senegal; US, United States, UY, Uruguay.

An analysis of recent FC1 phylogenetic structure (in both full genome and HA1 data sets) reveals two closely related subclades at this point in time: the viruses circulating in the US, and those collected from the 2019 European outbreak. The directions of transmission appear complex, but since the virus in the United States was circulating for a few years prior to 2019, that was likely transferred from the US to the United Kingdom and elsewhere in Europe. A few viruses collected in California in 2019 clustered with the European subclade, noted in horses that had been imported from Europe (metadata of samples collected by UC-Davis for this study). Some 2020-2021 US cases in the Northeastern states of New York and Rhode Island also more closely match the UK samples, suggesting likely transfers from Europe back to the USA. Further sampling in the USA, particularly from the Northeast region, is needed to clarify the relatedness of the North American and European FC1 viruses at this time.

### FC2 represents an under sampled and likely limited clade of EIV with current circulation in some regions of Asia

The public repositories have limited numbers of full genomes from the FC2 clade, although more HA1 sequences are available. Sampling appears to be relatively sparce, with multiple samples and sequences from notable outbreaks, so that there is extensive genetic redundancy among the samples available. Our FC2 full genome ML tree showed no variation from the full H3N8 data set analysis (Figure 5A, Figure 1). The more substantial HA1 data set (n=254, Figure 5B) clearly revealed geographic distribution of recent FC2 samples: while FC2 were centered in Europe between around 2000 and 2014, the clade appears to have been transferred to Asia around 2009. Europe has seen fewer FC2 cases, with last reports in 2017, while since 2018 only a single subclade has been present in mainland Asia. The small number of full FC2 genomes (Figure S1G-H) resulted in Bayesian analyses characterized by high statistical uncertainty (Figures S2A-B, S9-11).

**Figure 5.**
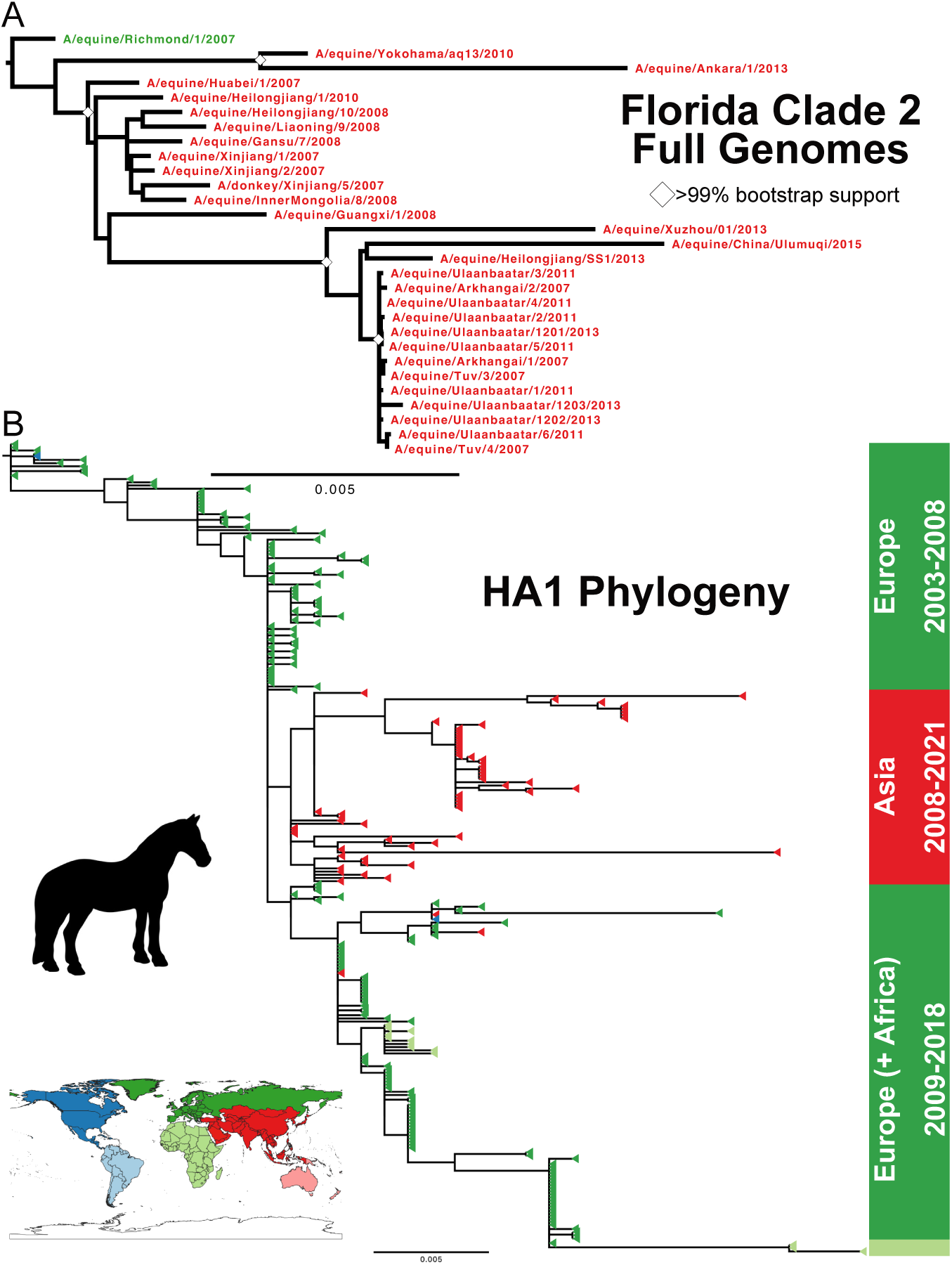
Evolution of the H3N8 EIV clade FC2. (A) ML tree of full genome data set of FC2, with taxa color defined by worldwide region. Tree was rooted to earliest full genome sample, Richmond/1/2007. Scale denotes nucleotide divergence. White diamonds at major nodes represent bootstrap support >99%. (B) ML tree of the expanded HA1 data set (n=254), also colored by regional origin. Tree was rooted to earliest HA1 FC2 sample, NewMarket/5/2003. Dates of and geography of regional subclades show that FC2 is now centered in mainland Asia.

We also note that FC2 isolates have been identified in various equid species (Mongolian horses, Arabian horses, Donkeys), as well as dogs, although their epidemiological relevance is unknown. Horses in Asia appear to be regularly exposed to avian influenza but have generally failed to support a new emergent strain due to a lack of adaptation (80).

### H3N8 clades show distinct mutational signatures reflecting host biology and unique epidemiology

We next compared H3N8 evolution in different hosts to see if there were obvious signatures of host-specific evolution after transfer to, and sustained transmission in, dogs. The rate of genome evolution (i.e. nucleotide substitutions per site per year for each segment) for each clade showed that substitution rates in CIV and FC1 were essentially the same in all segments except HA (Figure 6A), suggesting that if new-host adaptation was present it was masked in the general background rate of genetic drift (81, 82). In contrast, the HA, NA, and M1 substitution rates are significantly greater within the FC1 clade than the FC2. All H3N8 clades within horses and dogs exhibit lower substitution rates than reported for influenza circulating in human host populations (huH3N2, Figure 6A), particularly in HA and NA where antigenic selection would be most apparent.

**Figure 6.**
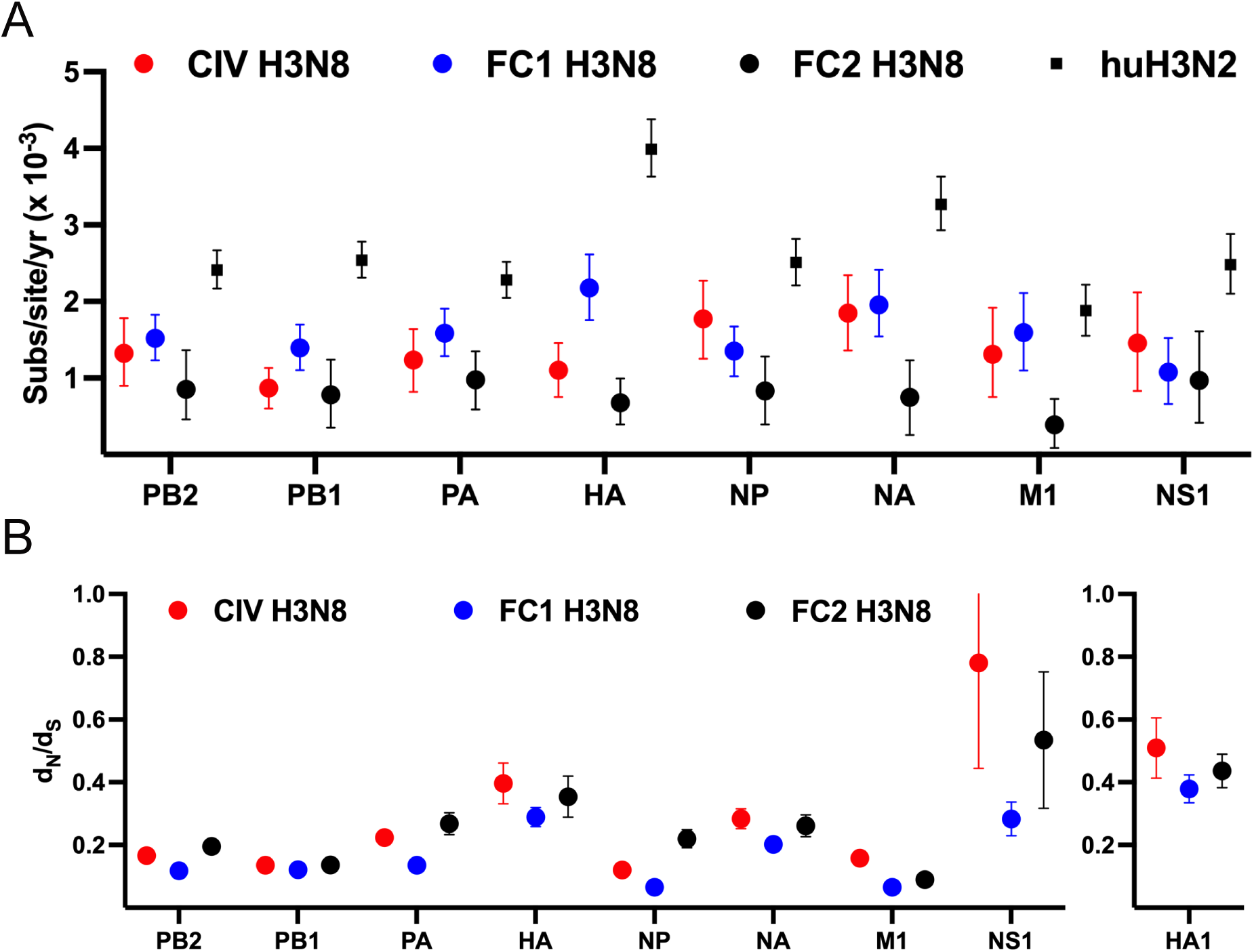
Molecular evolution of the H3N8 clades. (A) The nucleotide substitution rate for each segment major ORF in each clade are compared to known huH3N2 reference data set (102). Datapoints are mean values and error bars represent 95% highest posterior density (HPD) values. (B) The mean dN/dS for each segment major ORF was determined using the SLAC program from the Datamonkey platform. Datapoints are mean values and error bars represent 95% confidence intervals (CI).

We have previously reported comparative analyses of CIV and FC1 through 2015 (83). Only two CIV isolates have been generated since this time, while FC1 has continued to circulate widely in different regions of the world. Here, to better identify any residues subject to natural selection, we have updated that analysis based on the many additional the full genome sequences of the different clades. To do this we utilized several approaches in the Datamonkey analytical toolset (https://datamonkey.org/, (84)). Using SLAC (Single-Likelihood Ancestor Counting) we measured the mean dN/dS ratio of each segment for each clade (Figure 6B). An elevated ratio is seen in the HA ORF (particularly in the HA1 antigenic domain), but with no significant difference between clades. The divergent selection acting on the CIV and EIV H3N8 NS segment, reflected in variant segment length, has previously been analyzed in detail (85). In sum, it appears that FC1 is undergoing greater evolutionary change in horses than the FC2 clade, or than the H3N8 CIV in dogs up until 2016.

### CIV early mutations suggest host adaptation to dogs, yet with no change in epidemiology and avoidance of extinction

The sequences of viruses spilling over into new hosts will reflect the composition of the single virus that makes the initial transfer, and then will likely undergo adaptation to the properties and physiology of the new host animal, including optimizing onward transmission (3). We therefore compared the sequences and evolution of the viruses that were closely related to the common ancestor of CIV (preC), the evolution of H3N8 CIV in its new host, and parallel continued evolution of the early FC1 H3N8 EIV in horses (Figure 7A). This revealed multiple transitional mutations in CIV (Figure 7B), some of which have been previously analyzed (42, 83, 85, 86). However, these genetic changes appear not to have allowed CIV to increase its spread among the general dog populations, such that the virus continued to be maintained as shelter-driven outbreaks which resulted in viral extinction, likely through stochastic fade out (46). It is also likely that selection on the virus for spread within dense and high turn-over shelter or kennel populations did not result in the selection of mutations enabling enhanced spread in the household dog population required to develop a true epidemic or pandemic (45). Despite this successful transfer of the H3N8 EIV to dogs, all other spillovers have resulted in only single canine infections or short-lived local outbreaks (47).

**Figure 7.**
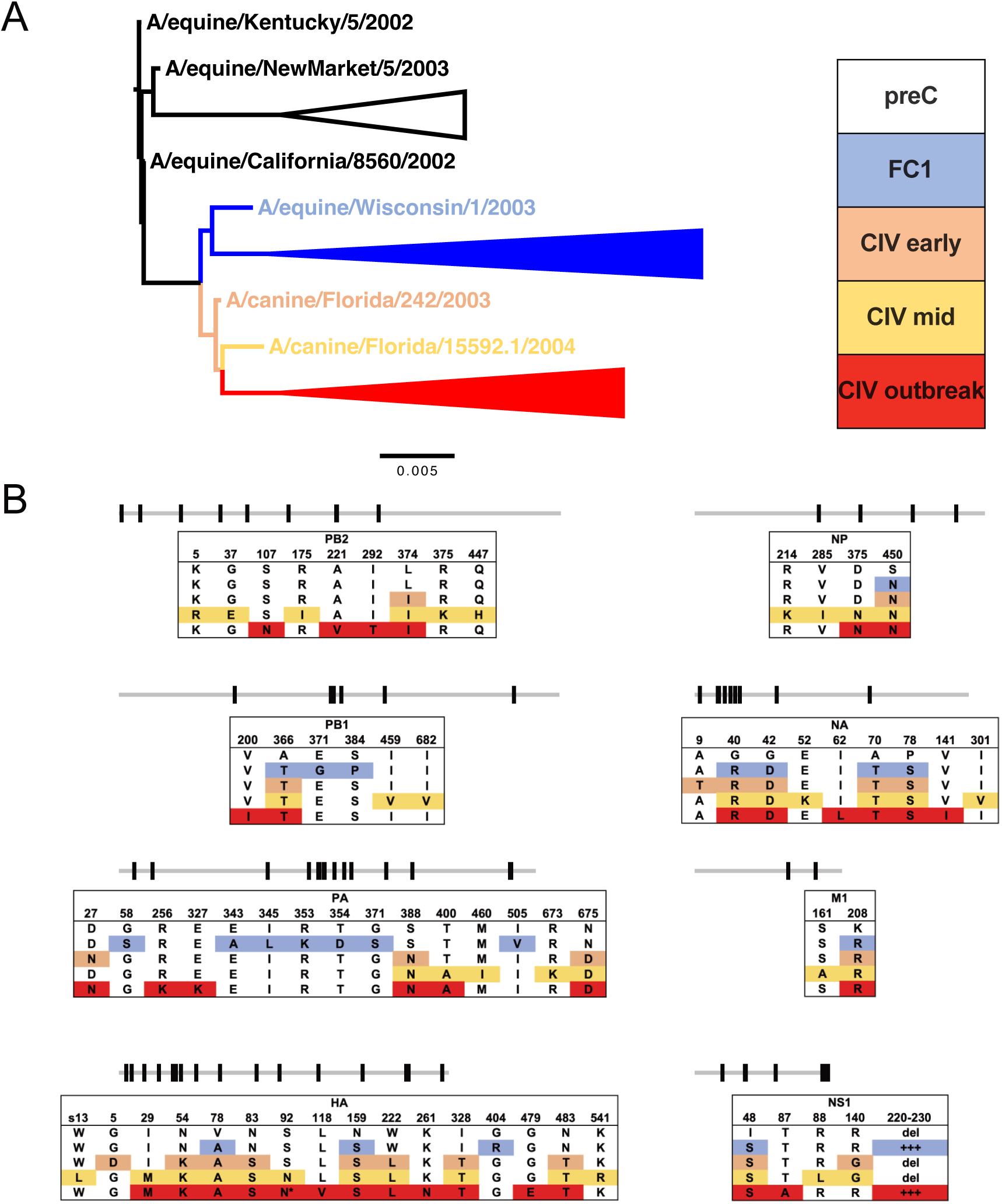
Early molecular evolution of H3N8 clade divergence and CIV emergence. (A) The H3N8 whole genome ML tree focused on the earliest preC, FC1, and CIV taxa for comparative alignment. (B) Key nonsynonymous changes in the earliest diverging taxa are identified by genome segment and position. Some late CIV mutations (*) are not present in all outbreak subclades.

It is also noteworthy that the H3N8 EIV FC1 acquired additional mutations during its parallel spread in horses. There is some evidence that H3N8 may have undergone further host adaptation in horses over time (87). This change in evolutionary potential of the EIV may suggest that the preC consensus genome was particularly suited to clade divergence and host-switching ∼1999. While some antigenic selection may occur (22, 40, 88), most mutations are not in the antigenic sites of the HA or NA, and likely represent genetic drift during the outbreak-driven circulation of FC1 (Figure 4). Discerning true antigenic evolution from background genetic drift will require greater surveillance and testing of immunity in horses.

## Discussion

A detailed understanding of the replication and spread of infectious diseases allows effective control strategies to be devised. While endemic human influenza outbreaks appear to involve common global movement and widespread infections, IAVs in other hosts involves more complex epidemiology, often with constrained geographic spread, and many outbreaks dying out without obvious human interventions. The recent apparent extinction of the Influenza B Yamagata lineage in humans suggest that influenza viruses may have a more tenuous foothold in mammalian host population than has sometimes been assumed (89).

### Equine and canine H3N8 have different evolutionary fates

The analysis presented here focuses on the avian-mammalian emergent H3N8 viruses: first in their original equine hosts in which they have been circulating for 60 years, and then with comparisons to the same virus in dogs, to understand the divergent evolution of the same virus in different hosts. While EIV continues to circulate and infect horses around the world, we report here that CIV was last seen in 2016 in the Northeast United States, and is apparently now extinct.

H3N8 CIV was likely entirely contained to the United States during its ∼17 years of transmission in dogs, as no cases of this lineage have been reported in other countries. Our new complete genome sequencing of the H3N8 CIV isolates from throughout the course epidemic show that its geographic structure arose from an initial burst of widespread disease that may have been associated with the movement of infected greyhounds. After 2007, only local transmission was seen in Colorado and close by in Wyoming, up to around 2010 (90). In the Northeast of the USA the virus was first reported in New York in 2005 and in Pennsylvania in 2007. The virus continued to circulate in New York, where it was isolated from short-lived outbreaks that occurred after introduction of virus (likely due to transport of infected dogs from the shelters where the virus was endemic), and those were seen in Connecticut, Pennsylvania, Vermont, Maine, and Massachusetts up until 2016 (45, 46). Our analysis confirms that the outbreaks in Colorado and the Northeast were distinct, and also that after 2012 only a single New York-centered lineage of virus continuing to circulate in dogs.

### Equine influenza virus circulates widely in the USA and in different regions of the world

In contrast, the H3N8 EIV has been found throughout North America for the 60 years since it was first isolated. Ongoing passive surveillance systems confirm US circulation, with some observed seasonality (26). However, frequent international transfers of the virus are also a feature of its epidemiology. We confirm here that outbreaks in several countries around the world were likely directly, or indirectly, caused by viruses from North America. Those included widespread outbreaks in South America that started in 2018, and in the United Kingdom and elsewhere in Europe starting in 2019. Similarly, viruses from Japan, Australia, Malaysia have been closely related to viruses from the USA. Recent outbreaks in Western Africa also fit this pattern, as viruses from Senegal, Nigeria, and Niger were closely related to those from South America, which were due to viruses originating in the USA (76). SARS-CoV-2 drastically changed human behavior during 2020 and 2021, with international travel spending being reduced by 76% in 2020 compared to 2019 following international travel bans imposed by many countries (91). Equine sport activities in the United States and around the world were consequently impacted by varied economic curtailment, including the international transfer of horses. Yet we see continued FC1 infections in the United States between 2020 and 2022, and a potential inter-relatedness between US and European samples in that time, suggesting a potential return to the norm. Overall, it appears that the FC1 lineage of EIV is maintained in the horse population of the United States, and that provides viruses causing outbreaks in other regions of the world (77).

### What host differences explain the different viral evolutionary fates?

As dogs usually live in households as single animals or in small groups as their population structure and movement patterns likely differ from that of horses. Larger numbers of susceptible dogs may be found in animal shelters, kennels, or dog day cares, and these will allow increased transmission. CIVs will spread rapidly within most shelters, and then the virus will die out once the population has all been infected and become immune, or transmission may be stopped within the shelter by quarantining affected dogs (45, 46). Horses are often found in groups or stables, yet its impact on transmission is unclear. The transfer of horses for sport (including internationally) is likely a large driver of EIV dissemination. Recent estimations by USDA-APHIS report that 57.8% of equine operations in the US transported any resident equid by vehicle off the operation and back in the preceding 12 months; 14.1% were >500 miles, 29.7% of operations transported equids to an adjacent state, and 11.8% transported equids beyond adjacent states; while only 0.7% of operations transferred horses internationally (28). Additionally, there appear to have been no international transfers of dogs infected with H3N8 CIV. Given the short lifespan of dogs, limited circulation of H3N8, and the absence of a licensed vaccine until 2009, we assume very little immune pressure on the CIV H3N8. Equids may be similar to humans as long-lived vertebrates where relatively high levels of anti-EIV antibody immunity are induced due to either infection or vaccination but wane in time between lifetime exposures (92).

Interestingly, the reservoir of H3N2 CIV in China and Korea appears to have been dogs being bred in large numbers, with infected dogs then transported to the USA or Canada to cause outbreaks among different dog populations. However, once again there has been no onward transmission of that H3N2 CIV to other countries from North America, suggesting that the recent US dog population structure may represent a ‘dead-end’ for endemic IAV circulation (51).

## Conclusion

Our analysis of H3N8 evolution in the horse and canine clades show similar levels of variation that are lower than that observed for influenza in humans. The data therefore suggest that host population structure and epidemiology rather than specific adaptive evolution are driving the sustained transmission of FC1 in horses. Given the extinction of CIV H3N8, this suggests that well targeted interventions could in turn result in extinction of EIV H3N8 FC1 in horses. This likely requires a more refined epidemiological understanding of circulation in the United States, updated vaccination formulation for FC1 and greater uptake. It may further require screening of international transfers of horses for influenza to reduce worldwide transmission, particularly the export of US-borne FC1 lineages, as well as the ‘return’ of viruses to the USA from Europe. Rational human intervention may be necessary to ‘break’ the host-structure epidemiology that is sustaining this virus.

## Materials and Methods

### H3N8 Sample Collection

Samples were obtained from various diagnostic centers following routine passive diagnostic testing for respiratory disease (listed in Table S1, red). Samples were received as either nasal or pharyngeal swab material, virus isolated in culture/egg passages, or extracted total nucleic acid (tNA) first utilized for quantitative reverse transcription-PCR (qRT-PCR) positivity. vRNA was extracted from clinical swab samples and isolated using the QIAamp Viral RNA Mini kit. Purified vRNA or tNA was then either directly used for influenza multi-segment RT-PCR or stored at -80°C.

### Generation of Influenza Virus Sequences

Genomes from samples were generated as cDNAs using a whole genome multi-segment RT-PCR protocol as described previously (93, 94). A common set of primers (5′ to 3′, uni12a, GTTACGCGCCAGCAAAAGCAGG; uni12b, GTTACGCGCCAGCGAAAGCAGG; uni13, GTTACGCGCCAGTAGAAACAAGG) that recognize the terminal sequences of the influenza A segments were used in a single reaction with SuperScript III OneStep RT-PCR with Platinum *Taq* DNA polymerase (Invitrogen). Following confirmation by gel electrophoresis, viral cDNA was purified either by standard PCR reaction desalting columns or with a 0.45X volume of AMPure XP beads (Beckman Coulter). Nextera XT libraries were generated with 1 ng of cDNA material, while Nextera FLEX libraries were generated with 150-200 ng of cDNA (Invitrogen). Libraries were multiplexed, pooled, sequenced using Ilumina MiSeq 2 X 250 sequencing, and consensus sequences were determined for each genome segment after assuring full genome coverage.

Consensus sequence editing was performed using Geneious Prime 2022.1.1. Paired reads were trimmed using BBduk script (https://jgi.doe.gov/data-and-tools/bbtools/bb-tools-user-guide/bbduk-guide/) and merged. Each sequence was assembled by mapping to a reference sequence of a previously annotated equine or canine H3N8 sequence. Consensus positions had read depth >300 and >75% identity. We did not observe unresolved positions under these conditions from the samples generated in this study.

### Phylogenetic Analysis

H3N8 nucleotide sequences were downloaded from the NCBI Influenza Virus Database and combined with the consensus genomes generated in this study (95). We focused on genomes from 2001 to present, encompassing the formation of the major virus clades (FC2, FC1, CIV). Both inter- and intra-subtype reassortants were identified using RDP4 (seven methods: RDP, GENECONV, Bootscan, MaxChi, Chimaera, SiScan, 3seq) and excluded if found to be a statistically significant outlier in two or more methods (96). Our total data set comprised 157 full genomes (preC: 3, FC2: 29, FC1: 82, CIV: 43), with an expanded 611 HA1 reading frames (preC: 17, FC2: 254, FC1: 252, CIV: 88) (Tables S1-2). Sequences were manually trimmed to their major open reading frames (PB2: 2280nt, PB1: 2274nt, PA: 2151nt, HA: 1695-1701nt, NP: 1497nt, NA: 1410-1413nt, M1: 759nt, and NS1: 654-693nt) and either analyzed separately or concatenated with all other genome segments from the same virus sample. HA1 nucleotide sequences were trimmed to cover amino acids 1-329 (H3 numbering). Nucleotide sequences were aligned using MUSCLE (97). Maximum likelihood (ML) phylogenetic analysis was performed by PhyML (98), employing a generalized time-reversable (GTR) substitution model, gamma-distributed (Γ4) rate variation among sites, and bootstrap resampling. Both MUSCLE and PhyML were run under the Geneious Prime platform. ML trees were visualized and annotated using FigTree v1.4.4 (http://tree.bio.ed.ac.uk/software/figtree/).

The temporal signal (i.e. molecular clock-like structure) of the data was assessed by a regression of root-to-tip genetic distance against date of sampling using our ML tree and the TempEst v.1.5.3 software (99). Accurate collection dating to the day (YYYY-MM-DD) was utilized for all samples where this information was available. The sampling dates used for all isolates are listed in Tables S1 and S2.

### Bayesian Phylogeny and Analysis

We used a Bayesian Markov chain Monte Carlo (MCMC) approach to better estimate virus divergence times and population dynamics. Analyses were performed in BEAST v.1.10.4 (100), where MCMC sampling was performed with a strict clock, a generalized time-reversable (GTR) substitution model with gamma distribution in four categories (Γ4), and assuming a Bayesian Skyline Plot (BSP) prior demographic model. Analyses were performed for a minimum of 100 million runs, with duplicates combined in Log Combiner v1.10.4. All output results were examined for statistical convergence in Tracer v1.7.2 (effective sample size [ESS] ≥200, consistent traces, removed burn-in at 10-15%) (101). Maximum clade credibility (MCC) phylogenetic trees were generated by Tree Annotator v1.10.4, and visualized by FigTree v1.4.4.

### Analysis of Selection Pressures

The relative numbers of synonymous (dS) and nonsynonymous (dN) nucleotide substitutions per site in each ORF of each segment were analyzed for the signature of positive selection (adaptive evolution, p<0.05 or >0.95 posterior) using a variety of method packages available in the Datamonkey package (https://datamonkey.org/, (84)): BUSTED (**B**ranch-site **U**nrestricted **S**tatistical **T**est for **E**pisodic **D**iversification), FEL (**F**ixed **E**ffects **L**ikelihood), FUBAR (**F**ast, **U**nconstrained **B**ayesian **A**pp**R**oximation), MEME (**M**ixed **E**ffects **M**odel of **E**volution), and SLAC (**S**ingle-**L**ikelihood **A**ncestor **C**ounting).

### Graphing and Statistical Analysis

All figure graphs were generated in GraphPad Prism v.9.5.0.

### Data Availability

Virus genome consensus sequences have been submitted to GenBank, and accession numbers of the sequences used (and generated) for this study are provided in Tables S1 and S2.

## Acknowledgements

We thank the sources of the new samples used in this study, including diagnostic labs and individual private veterinarians. We wish to thank members of the Parrish Lab for critical feedback and support on this manuscript, particularly Wendy Weichert. SER and TMC were supported by a project of the Kentucky Agricultural Experiment Station, Project No. KY-014067. PRM was supported by the Medical Research Council of the United Kingdom (Grant MC_UU_12014/9), the Horserace Betting Levy Board (Grants 797), the Biotechnology and Biological Sciences Research Council (Grants BB/V002821/1 and BB/V004697/1). CRP (and by extension BRW, ER, and IEHV) were supported by National Institutes of Health grant R01-GM080533. This work was partially funded by the Influenza Division of the Centers for Disease Control, Animal-Human Interface program, under contract #75D30121P12812 to CRP and LBG.

