## Supplementary material for "Understanding the Divergent Evolution and Epidemiology of H3N8 Influenza Viruses in Dogs and Horses": SuppFig (from another version of this preprint: version 1, 2023-03-23)

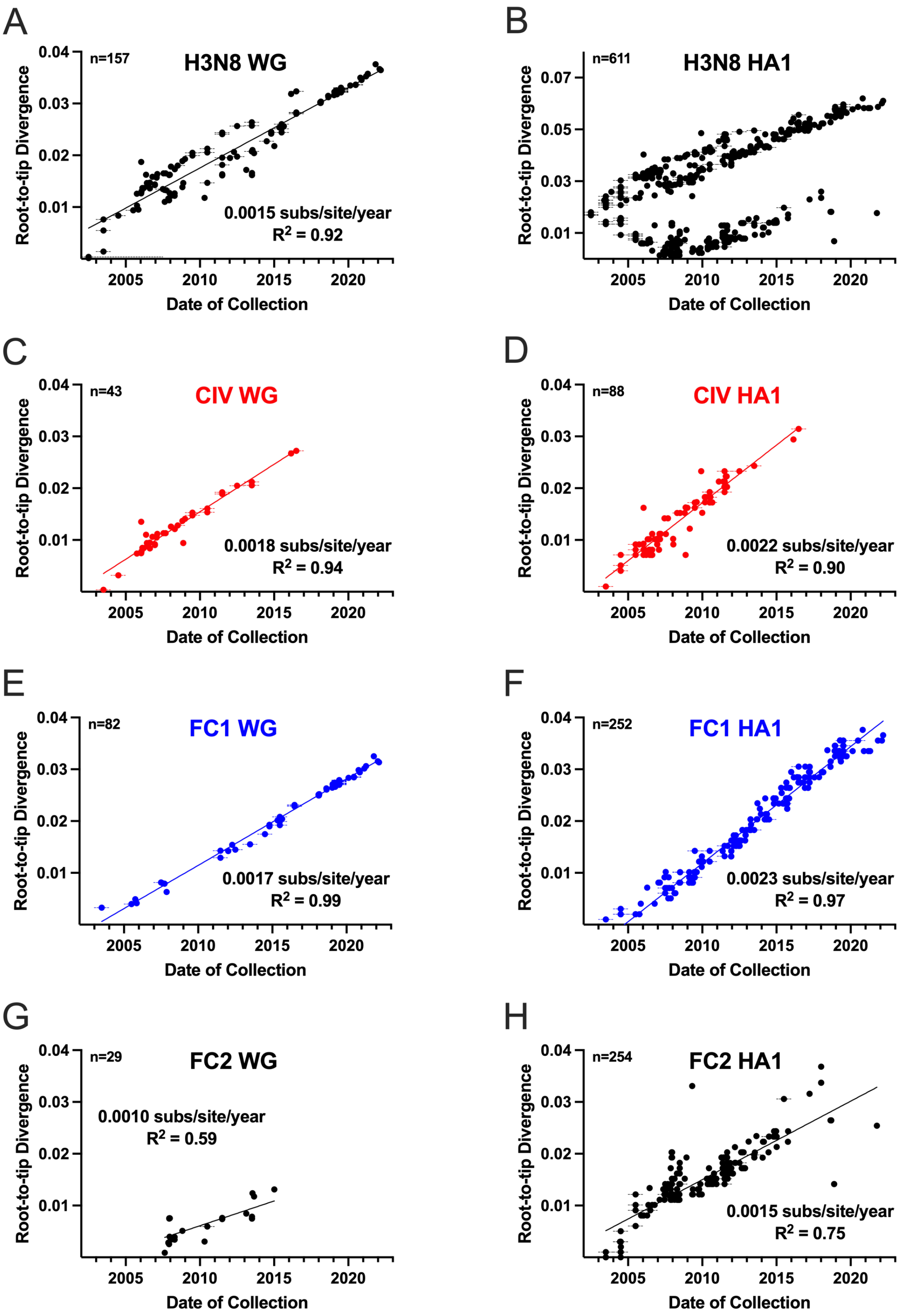


**Figure S1**. Temporal structure of H3N8 full genome and HA1 datasets analyzed using root-to-tip regressions (TempEst). Date of collection is to best accuracy (YYYY-MM-DD) with uncertainty denoted as horizontal error. Linear regression analysis is represented by the line, with output details provided on graph. The distinction between FC1/CIV and FC2H3N8 is apparent in the HA1 dataset (B).

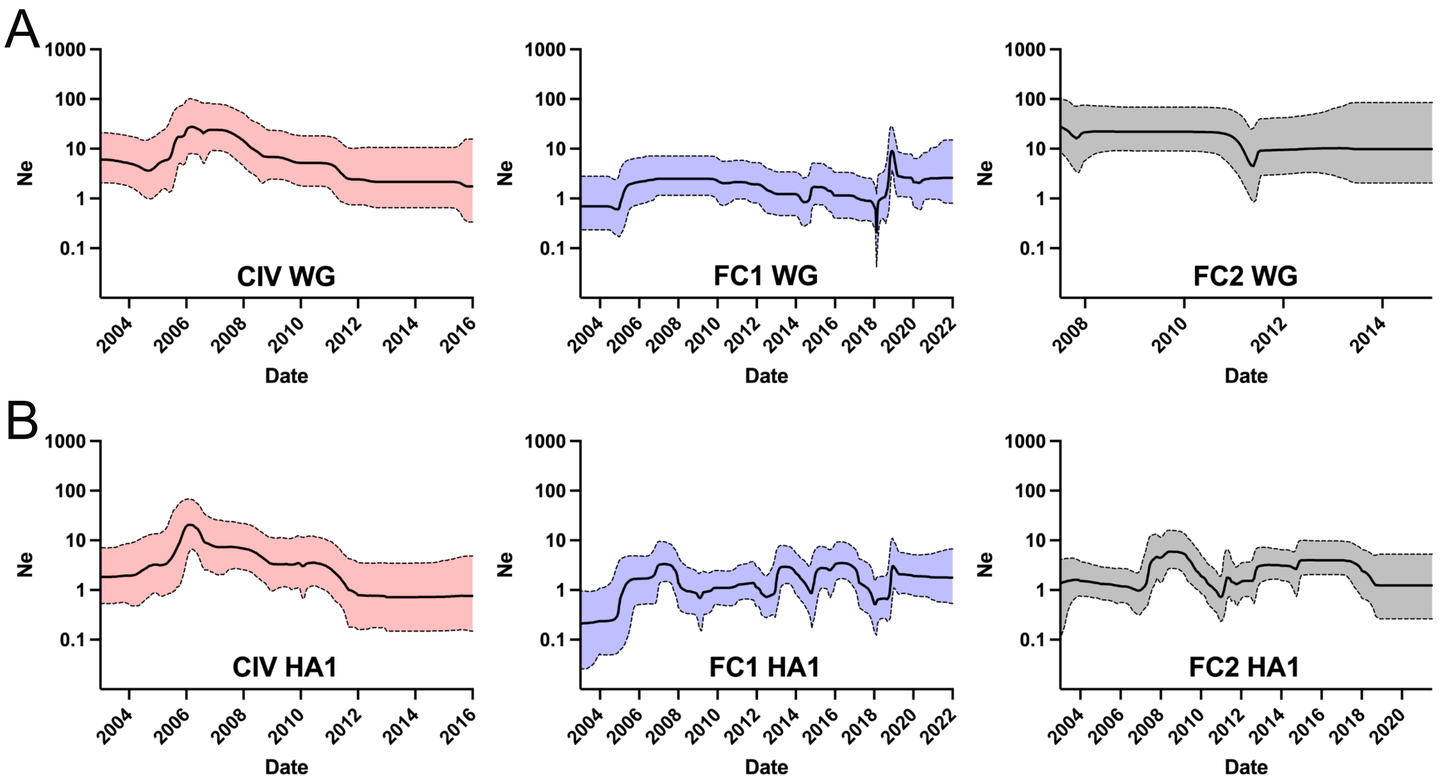


**Figure S2**. Bayesian Skyline Plots (BSP) of major data sets. Solid line is median value of effective population size (Ne). Shaded area between dotted lines are 95% highest posterior density (HPD) values. (A) Whole genome data sets. (B) HA1 data sets.

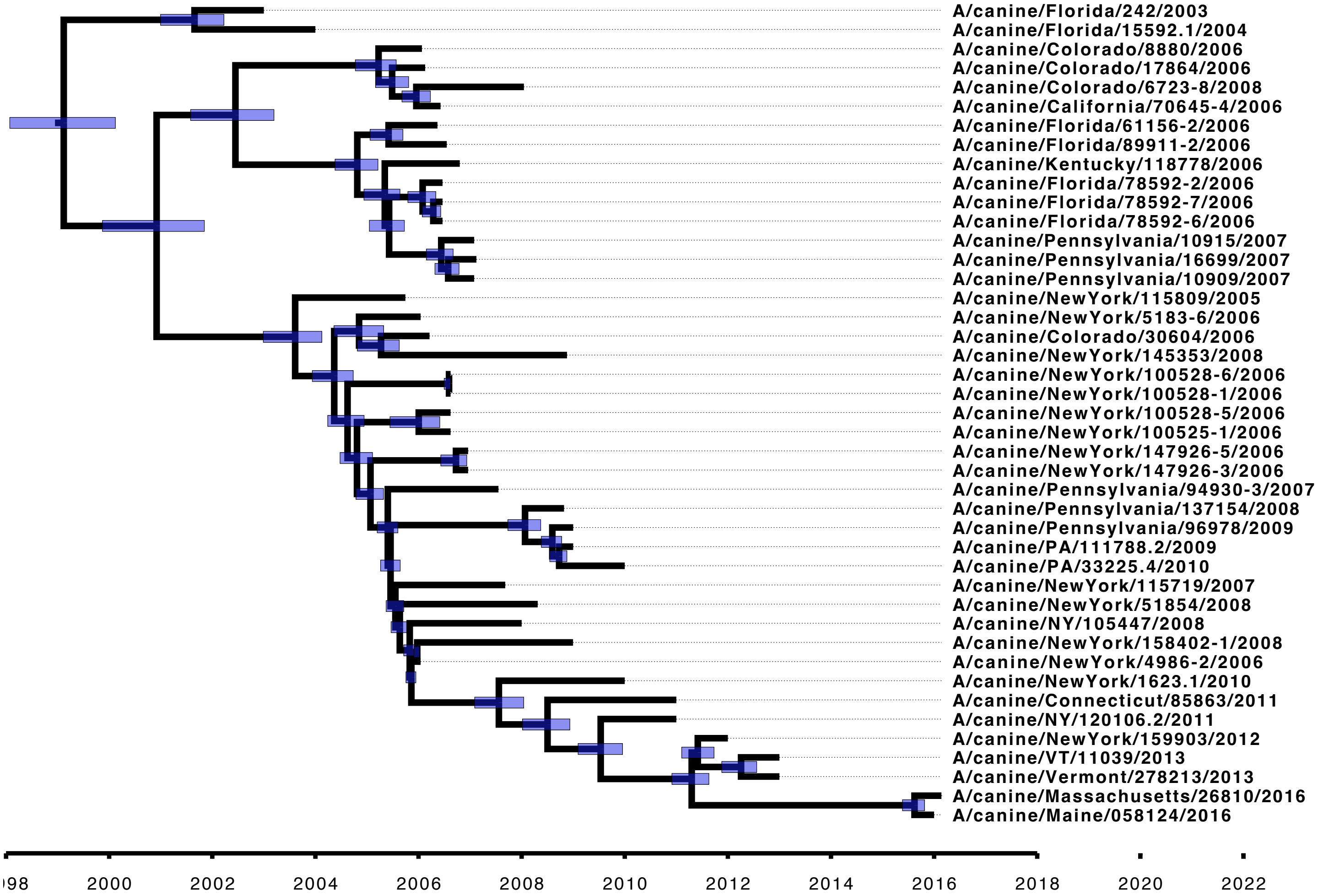


**Figure S3**. MCC tree of CIV full genome data set (n=43) estimated under a strict clock, Bayesian skyline prior. Tree is unrooted and scaled to dating by the youngest tip. Blue bars designate node heights (+/- 95% highest posterior density (HPD)).

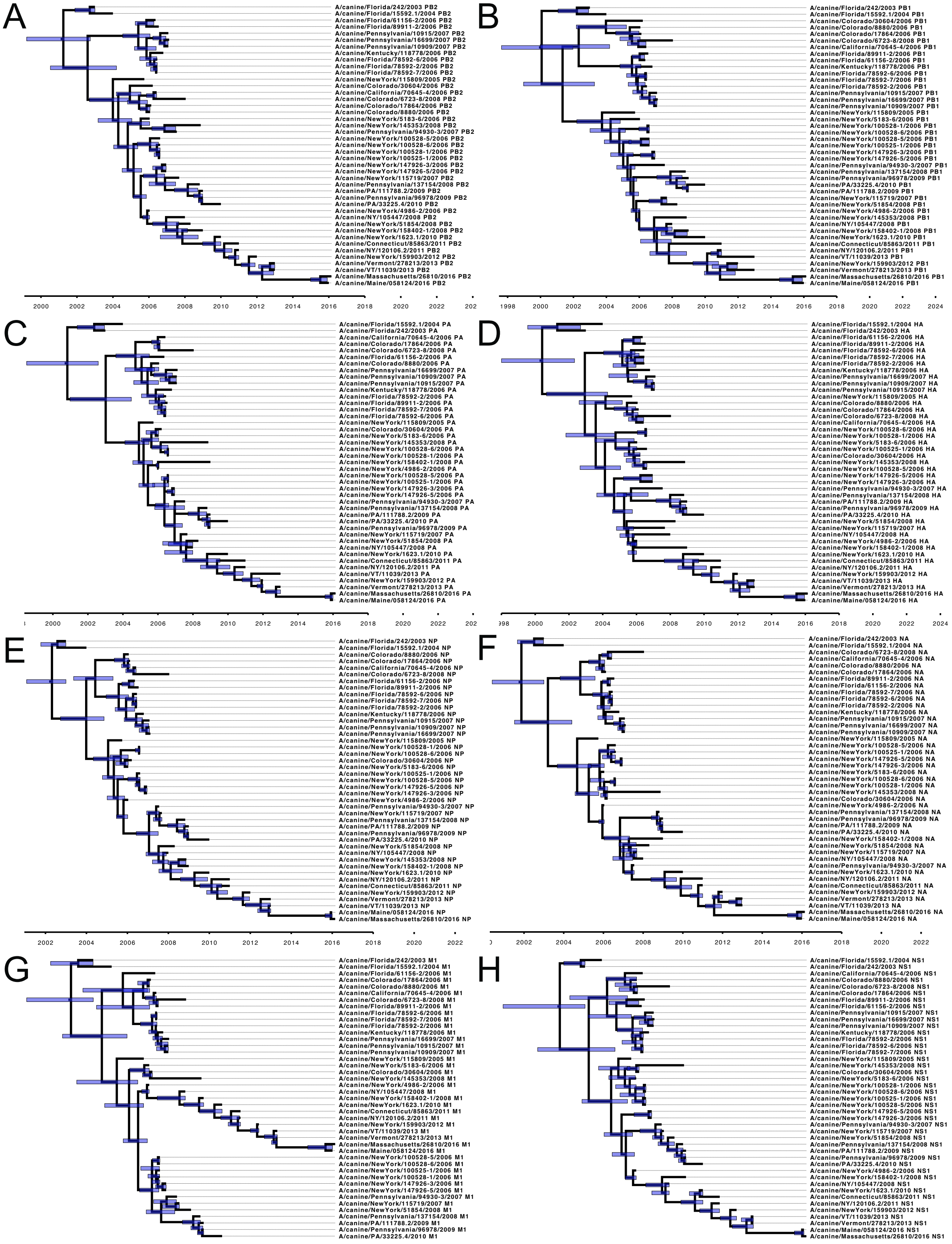


**Figure S4**. MCC trees of the major ORF of individual CIV segments (n=43). Trees are unrooted and scaled to dating by the youngest tip. Blue bars designate node heights (+/- 95% highest posterior density (HPD)). ORFs include: PB2 (A), PB1 (B), PA (C), HA (D), NP (E), NA (F), M1 (G), NS1 (H).

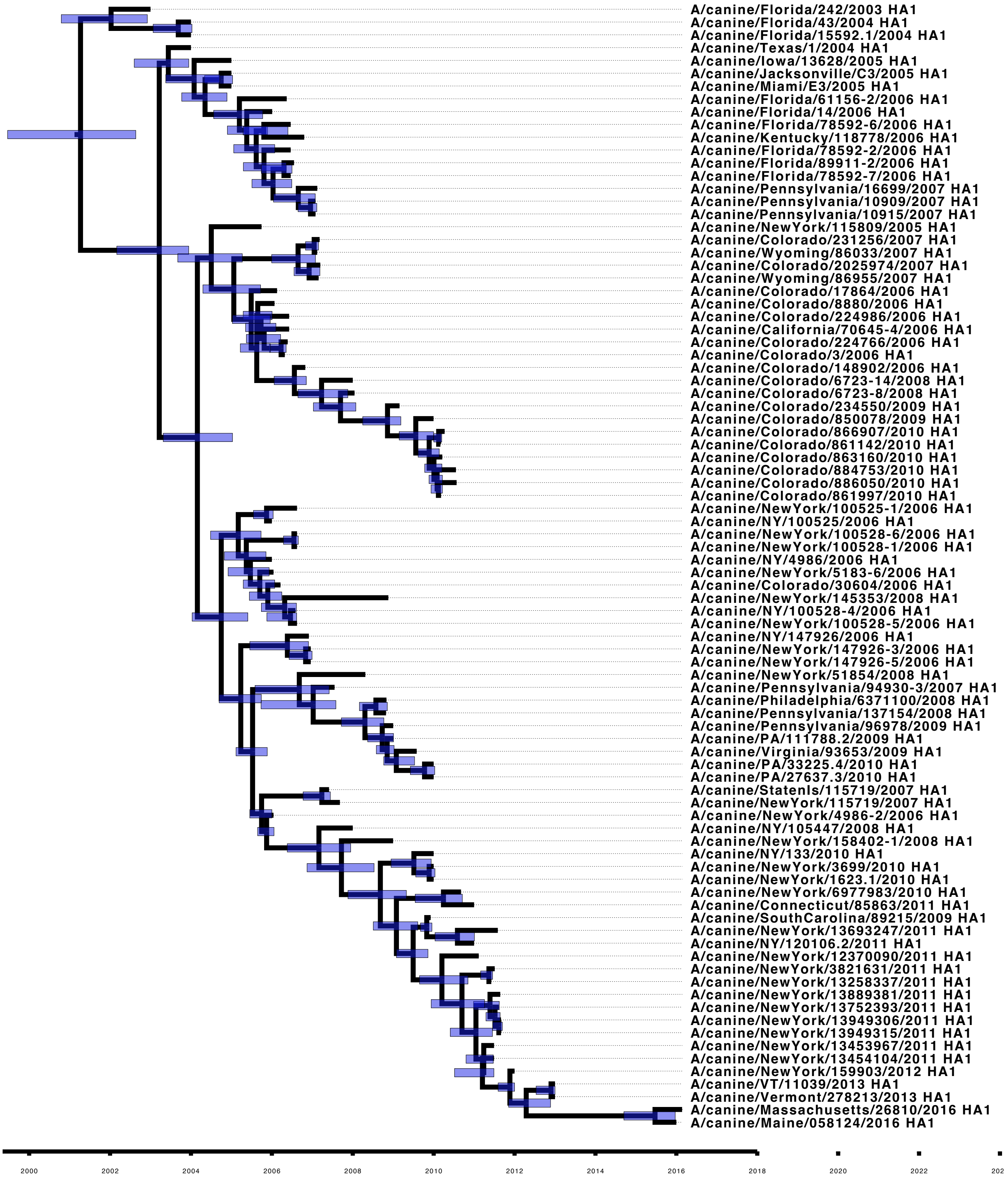


**Figure S5**. MCC tree of CIV HA1 dataset (n=88) performed under a strict clock, Bayesian skyline prior. Tree is unrooted and scaled to dating by the youngest tip. Blue bars designate node heights (+/- 95% highest posterior density (HPD)).

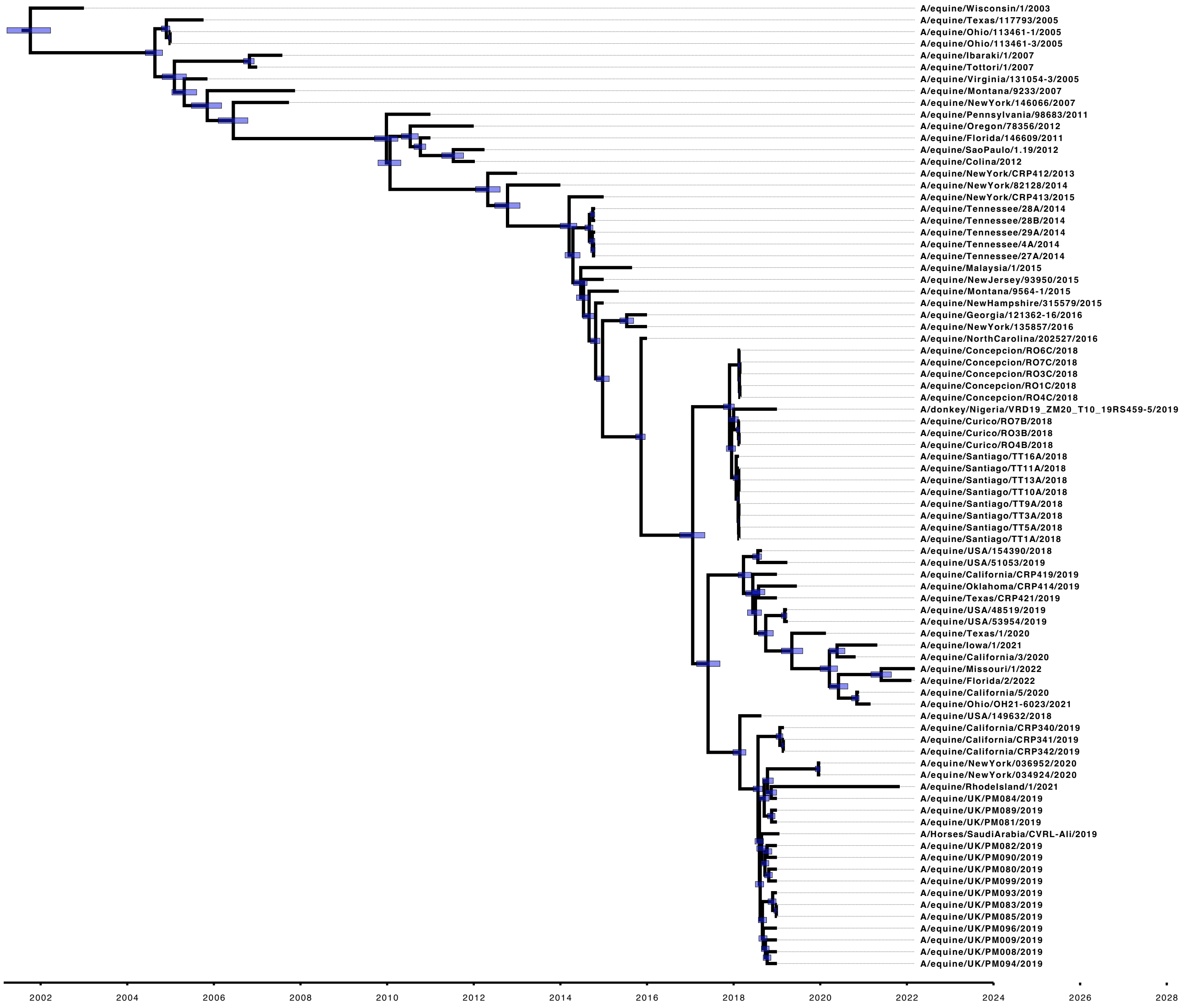


**Figure S6**. MCC tree of FC1 full genome dataset (n=82) performed under a strict clock, Bayesian skyline prior. Tree is unrooted and scaled to dating by the youngest tip. Blue bars designate node heights (+/- 95% highest posterior density (HPD)).

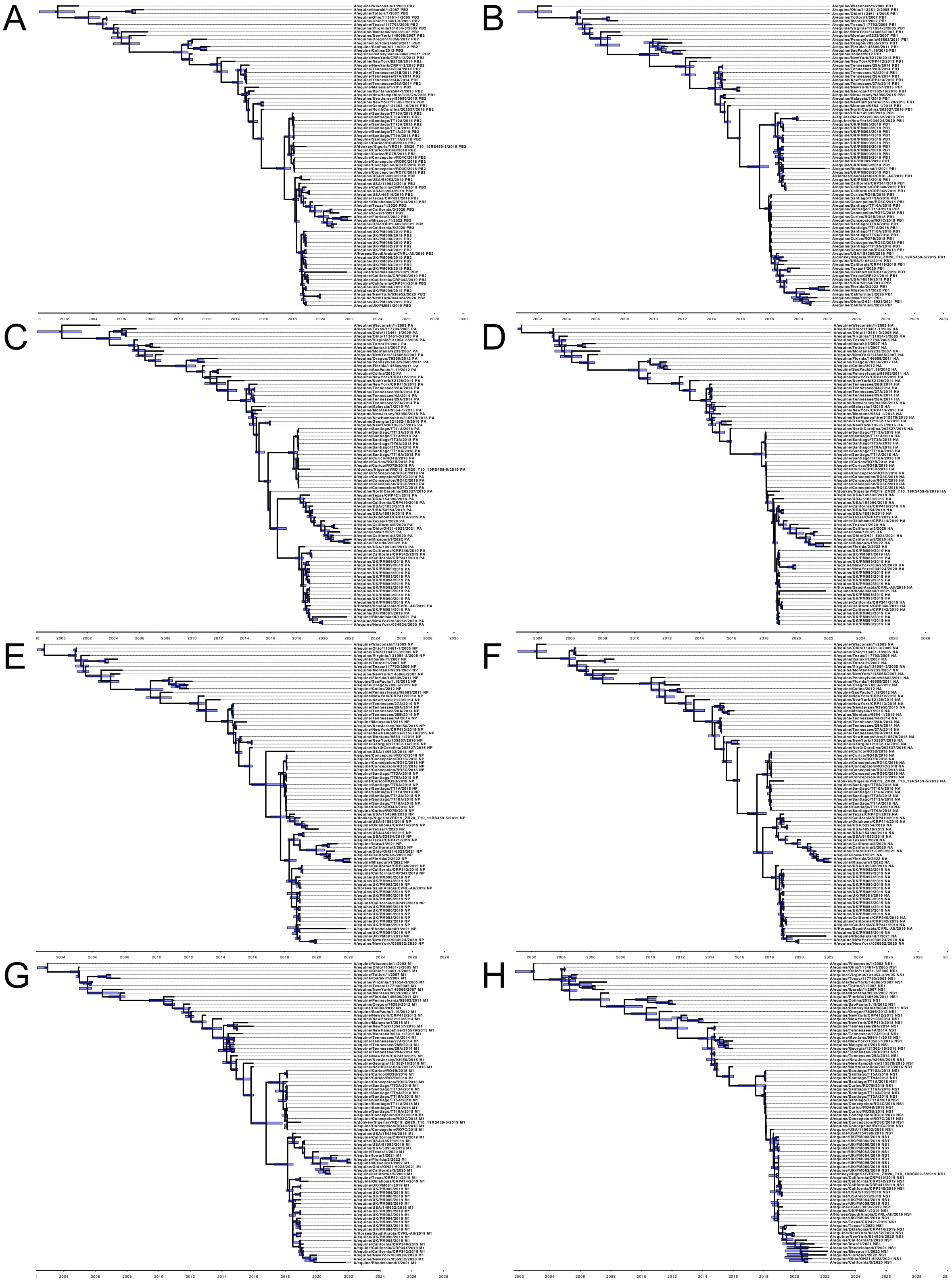


**Figure S7**. MCC trees of the major ORF of individual FC1 segments (n=82). Trees are unrooted and scaled to dating by the youngest tip. Blue bars designate node heights (+/- 95% highest posterior density (HPD)). ORFs include: PB2 (A), PB1 (B), PA (C), HA (D), NP (E), NA (F), M1 (G), NS1 (H).

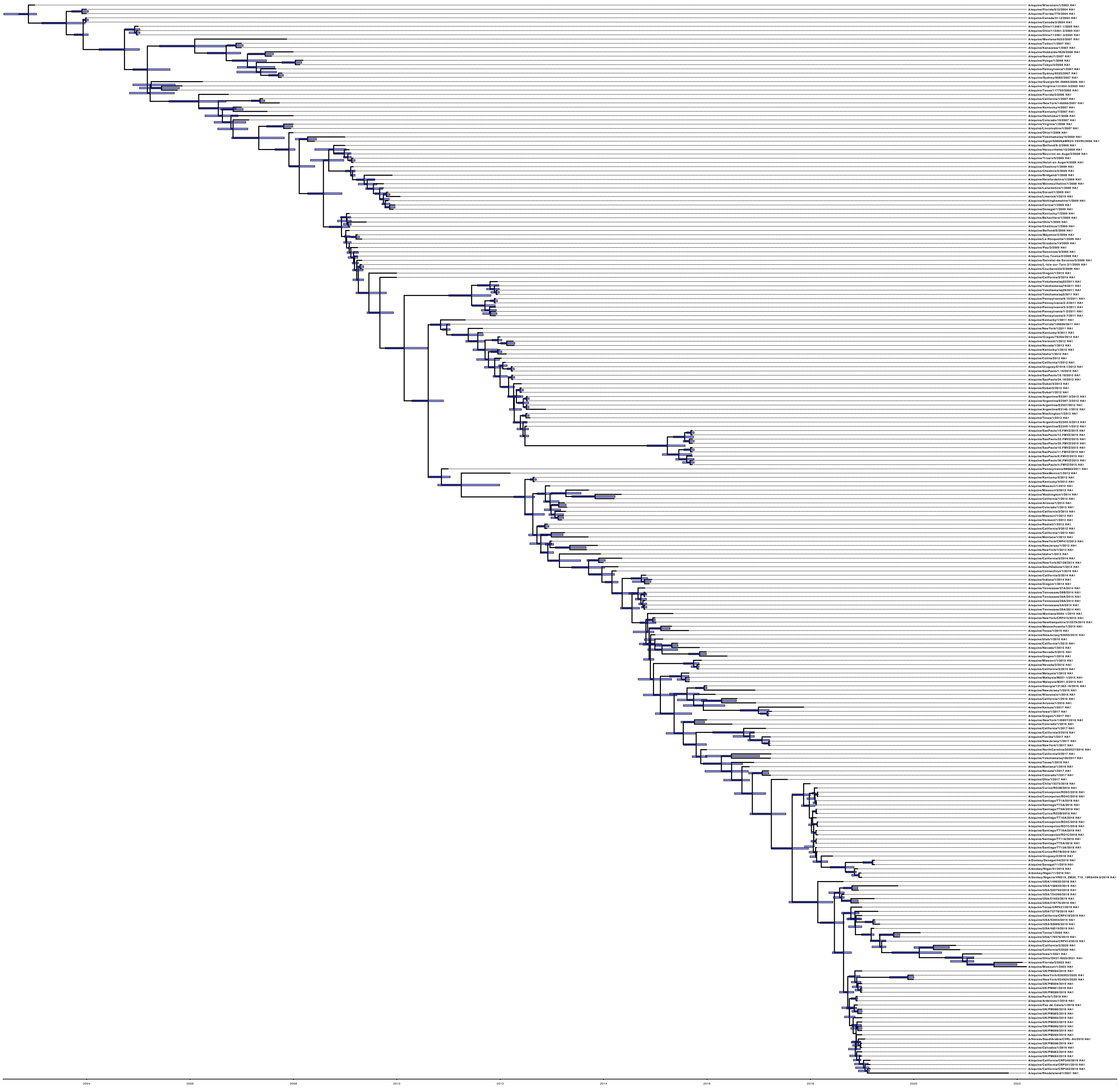

**Figure S8**. MCC tree of FC1 HA1 dataset (n=252) performed under a strict clock, Bayesian skyline prior. Tree is unrooted and scaled to dating by the youngest tip. Blue bars designate node heights (+/- 95% highest posterior density (HPD)).

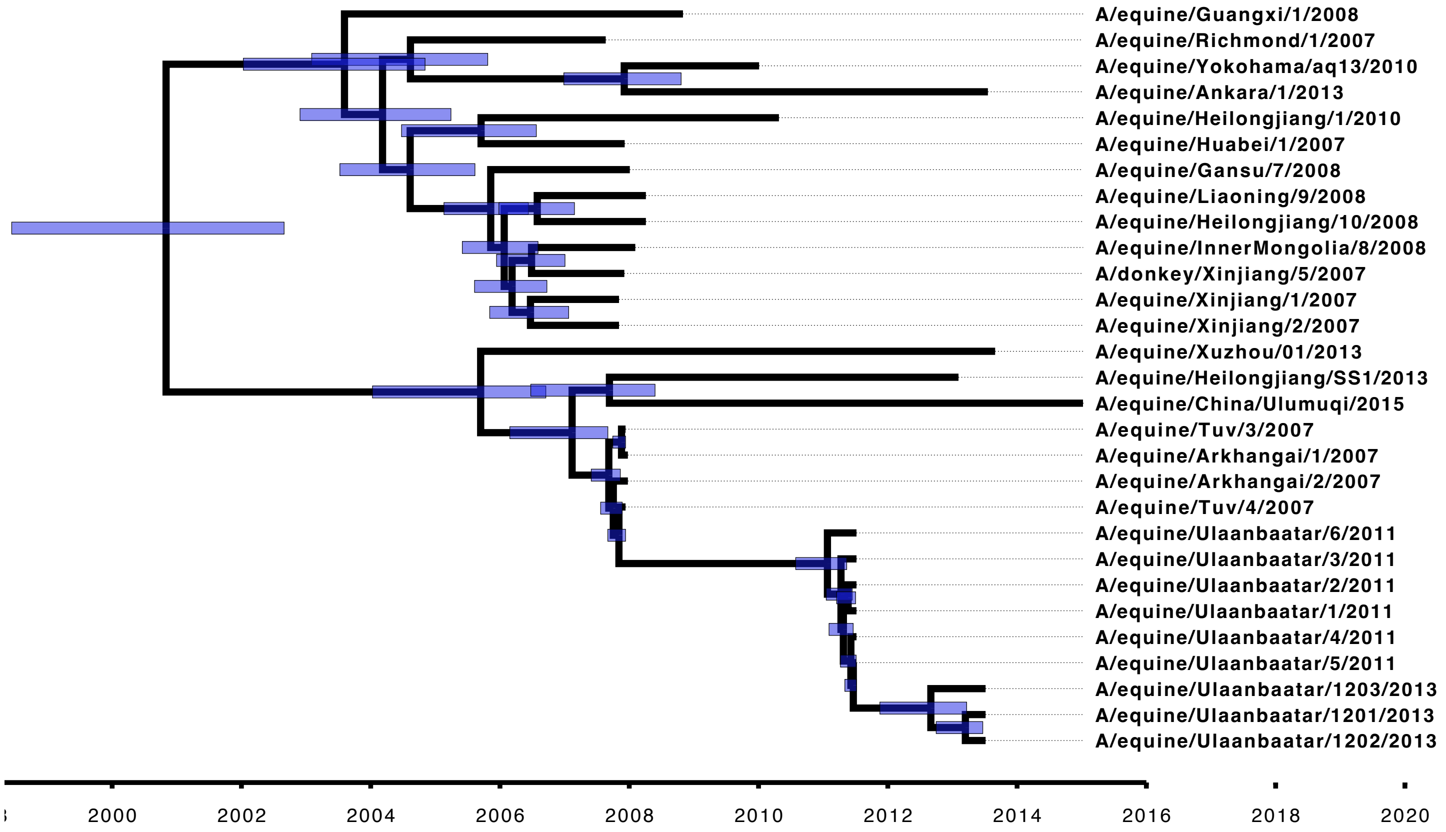


**Figure S9**. MCC tree of FC2 full genome dataset performed under a strict clock, Bayesian skyline prior. Trees are unrooted and scaled to dating by the youngest tip. Blue bars designate node heights (+/- 95% highest posterior density (HPD)).

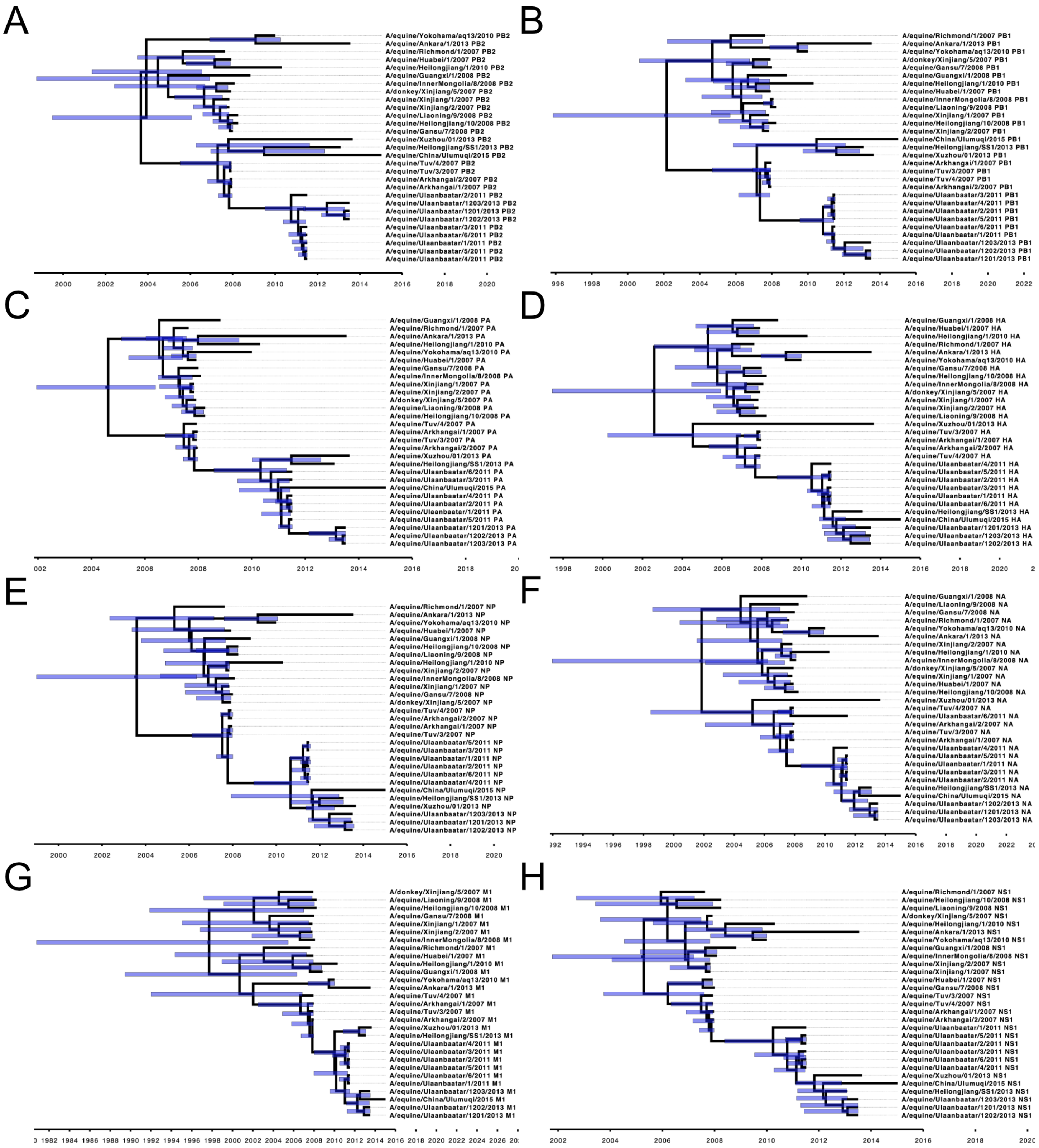


**Figure S10**. MCC trees of the major ORF of individual FC2 segments (n=29). Trees are unrooted and scaled to dating by the youngest tip. Blue bars designate node heights (+/- 95% highest posterior density (HPD)). ORFs include: PB2 (A), PB1 (B), PA (C), HA (D), NP (E), NA (F), M1 (G), NS1 (H).

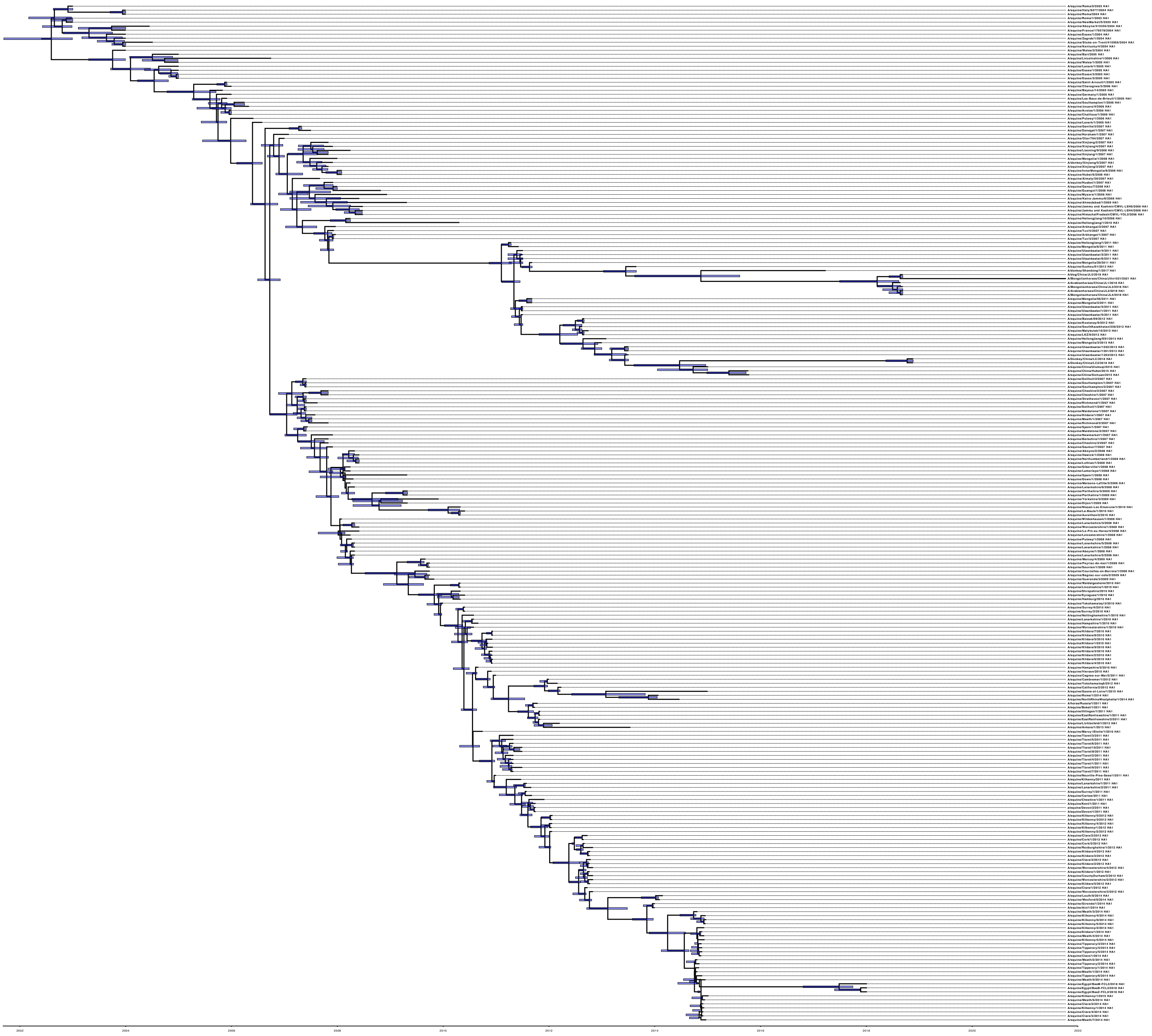

**Figure S11**. MCC tree of FC2 HA1 dataset (n=254) performed under a strict clock, Bayesian skyline prior. Tree is unrooted and scaled to dating by the youngest tip. Blue bars designate node heights (+/- 95% highest posterior density (HPD)).
